# Leveraging Comparative Phylogenetics for Evolutionary Medicine: Applications to Comparative Oncology

**DOI:** 10.1101/2025.02.11.637459

**Authors:** Walker J. Mellon, Beckett Sterner, J. Arvid Ågren, Orsolya Vincze, Matthew T. Marx, Stefania E. Kapsetaki, Ping-Han Huang, Bryan Yavari, Hunter W. McCollum, Barbara Natterson-Horowitz, Hannah Human, Cristina Baciu, Harley Richker, Diego Mallo, Carlo Maley, Luke J. Harmon, Zachary T. Compton

**Affiliations:** Arizona Cancer Evolution Center, The Biodesign Institute, Tempe, AZ; School of Life Sciences, Arizona State University, Tempe, AZ; Lerner Research Institute, Cleveland Clinic Foundation, Cleveland, OH; Institute of Aquatic Ecology, Centre for Ecological Research, 4026 Debrecen, Hungary; Evolutionary Ecology Group, Hungarian Department of Biology and Ecology, Babeş-Bolyai; Frederick University, School of Health Sciences, Department of Pharmacology, Nicosia, Cyprus; Hellenic Open University, Patras, Greece; School of Mathematical and Statistical Sciences, Arizona State University; Indiana University School of Medicine, Indianapolis, IN; Department of Human Evolutionary Biology, Harvard University, Cambridge, MA; Department of Biological Sciences, University of Idaho, Moscow, Idaho; University of Arizona Cancer Center, Tucson, AZ; University of Arizona College of Medicine, Tucson, AZ

## Abstract

Comparative phylogenetics provides a wealth of computational tools to understand evolutionary processes and their outcomes. Advances in these methodologies have occurred in parallel with a surge in cross-species genomic and phenotypic data. To date, however, the majority of published studies have focused on classical questions in evolutionary biology, such as speciation and the ecological drivers of trait evolution. Here, we argue that evolutionary medicine in general, and our understanding of the origin and diversification of disease traits in particular, would be greatly expanded by a wider integration of phylogenetic comparative methods (PCMs). We use comparative oncology – the study of cancer across the tree of life – as a case study to demonstrate the power of the approach and show that implementing PCMs can highlight the mode and tempo of the evolutionary changes in intrinsic, species-level disease vulnerabilities.

---

> “We must learn to treat comparative data with the same respect as we would experimental results.”
>
> — (Maynard Smith and Holliday 1979, p. vii)

## Introduction

Evolutionary medicine seeks to understand the ultimate origin and persistence of human disease within the context of the species’ evolutionary history and that of their relatives (1–3). A central explanatory concept in the field has been the idea of evolutionary mismatch (4,5), which hypothesizes that contemporary maladies are often rooted in the dramatic shift in environment humans have undergone on a timescale that natural selection cannot meaningfully act upon. In the last decade, a plethora of human diseases have been explored through this lens, including autoimmune disorders, diabetes, heart disease, and psychiatric disorders (1,2,6–14).

The evolutionary mismatch framework has been incredibly productive. To date, however, the majority of research in evolutionary medicine has focused on humans (*Homo sapiens)* and our immediate ancestors, with relatively little attention paid to the context of broader phylogeny. For example, early attempts to explain the variation in cancer risk across taxa heavily relied on a life history framework of disease risk among humans and our closest relatives (15–19). Given the diverse ecological backgrounds in which species evolve, variant strategies will dictate the amount of energetic investment into somatic maintenance and other disease relevant domains. Moreover, the application of evolutionary trait models can drive deeper testing of the hypotheses developed from a life history framework of disease vulnerability. Life history theory has provided a powerful toolkit for understanding the evolution of reproductive and survival strategies across different species. More recently, this framework has been applied to studies of cross-species cancer prevalence, and a growing body of research suggests that cancer susceptibility may be linked to reduced energetic investment in somatic maintenance, favoring a high-volume reproductive output instead, as suggested by fast life history theory (15,20–22). Yet, there is more to be learned about human maladies when we expand our focal point to examine disease across deeper evolutionary history. In recent years, the integration of phylogenetic comparative methods (henceforth, PCMs), the study of evolutionary relationships among species, has provided such avenues for investigating the evolutionary origins and diversification of disease risk.

The comparative method is a cornerstone of evolutionary biology (23). However, it was not until the 1980s that the development of PCMs made significant headway, due in equal parts to the development of robust statistical approaches and confidence in DNA-based phylogenies (24–27). (“The comparative method of 1950 was indistinguishable from the comparative method of 350 BC,” as Ridley 1983, p. 6 put it(28).) By now, the rigor of statistical and computational techniques within comparative phylogenetics have come to rival those of population genetics. The continuous growth in computational methods and the accumulation of vast amounts of genomic and phenotypic data across a wide range of species has made PCMs increasingly popular. While attempts to apply PCMs to questions in evolutionary medicine have increased, efforts to outline ‘best practices’ for PCMs in evolutionary medicine have been sparse (29,30).

In this review, we argue that the integration of PCMs in evolutionary medicine can greatly enhance our understanding of disease risk, which may inform novel strategies for prevention and treatment of disease. Drawing on recent advances in PCMs and evolutionary medicine, we highlight the key benefits of this integrative approach and provide examples of its applications in the study of comparative oncology – the study of cancer across the tree of life (31–34). We begin by outlining some of the basics of the comparative method and then apply it to a previously published dataset of cross-species cancer prevalence. We end by discussing the challenges and limitations of using PCMs in the context of human health, and provide a roadmap for its applications in broader evolutionary medicine topics.

### A primer on comparative phylogenetics

The foundation of modern models of continuous trait evolution rests with Brownian motion (discrete traits - traits with distinct states - are typically modeled with a model called the Markov k-state, or Mk, model). Models of Brownian motion originally developed from attempts to explain the outcome of processes that are seemingly the product of random interactions, specifically the random movement of particles suspended in a medium (fluid or gas). First described while observing the “irregular motion” of coal dust on the surface of alcohol by Jan Ingenhousz, the Brownian motion is actually named after the work of botanist Robert Brown, some decades later. Brownian motion is the basis of a large family of mathematical models that are popular due to their mathematical tractability and are suitable for modeling phenomena where small stochastic displacements add up to approximate Gaussian fluctuations. In comparative phylogenetics, data that has changed over time is plentiful, in the form of the average values of continuous evolutionary traits within populations or species. Many such traits are well-fit by Brownian motion. Naturally, early phylogenetic studies and research applied the Brownian model to the divergence of traits over time rather successfully. So it follows that this is often the basis of the null hypothesis in models of phylogenetic context (35). Although one possible mechanism for random change fitting a Brownian model is neutral “genetic drift”, there are other possible mechanisms for Brownian motion involving selection (36,37). In cases where Brownian motion is the most suitable model for a trait, it is characterized by a “random walk,” signifying a lack of directionality or identifiable trends in variation. Given that Brownian motion is one of the most commonly best-supported models, it provides a useful null hypothesis in investigating alternative modes of phenotypic evolutionary change (38). Building on the foundational model of Brownian motion, alternative evolutionary models have been developed over the past decades that are now able to incorporate various evolutionary forces, such as directional selection, adaptive radiation, and evolutionary constraints, to better fit empirical data (39–43). Crucially, the advancement and sophistication of the computational modeling abilities have also expanded the range of questions we can inquire about the evolutionary forces driving the extensive variation we observe in species’ traits.

### Phylogenetic Variance-Covariance Matrices

Phylogenetic variance covariance (VCV) matrices are essential to phylogenetic statistical models. In phylogenetic comparative methods, many models predict that the observed average values of continuous traits across species will follow a multivariate normal distribution. To account for phylogenetic relationships, a matrix is constructed by summing the shared branch lengths between species from their most recent common ancestor to the root, capturing evolutionary relatedness. This matrix is then scaled according to the assumptions of a given evolutionary model to generate the variance-covariance matrix, which describes the expected covariances among species based on their shared evolutionary history (44). The phylogenetic VCV can be utilized in various statistical methods to model trait relationships, applying it in evolutionary model fitting to capture expected trait covariances and in phylogenetic generalized least squares (PGLS) to model residual error and account for the non-independence of species for regression estimates.

### Analyzing Models of Phenotype Evolution

A prominent extension of Brownian motion, used widely in mathematical and physical frameworks, is the Ornstein–Uhlenbeck (OU) model, which describes trait evolution that reverts toward an optimal or mean value. George Eugene Uhlenbeck and Leonard Ornstein discovered the model in 1930 to describe the velocity of a Brownian particle undergoing friction (resistance to change)(45). OU has broad applications in physics, finance, and evolutionary biology. The OU model of evolution was adapted from models to explain random movements in a statistical framework to describe the evolution of continuous traits in a population over time (46). Evolutionary biologists commonly interpret this model as incorporating both stabilizing selection and genetic drift to explain how a trait may evolve in response to stochastic displacements around a center, e.g. the fitness optimum (47). An OU model includes a stochastic component, similar to Brownian motion, while the deterministic component models describes the pull toward a medial value due to stabilizing selection or any other restraining force.

Beyond the OU model, other evolutionary models further expand on Brownian motion to capture various aspects of evolution. For instance, the rate trend model is variation of the Brownian motion model with linear trends in traits or step variance (i.e. how fast a trait spreads around a mean) that describe the rate of change in continuous traits over time (40,48). These rates can be either positive (indicating increasing rates over time) or negative (indicating decreasing rates over time). This model can explain how evolutionary pressures and environmental changes influence or inhibit rates of trait change.

The mean trend model in phylogenetic trait evolution accounts for a directional change in traits over time, incorporating both a random walk component and a consistent drift. Unlike the Brownian Motion model, which describes evolution purely as a stochastic process, the mean trend model adds a “drift” parameter to capture the gradual increase or decrease in trait values across evolutionary time. This drift represents a directional bias, suggesting that traits may not only evolve randomly but may also follow a specific trend due to persistent selective pressures or long-term directional processes (39,40).

The white noise model represents a scenario in which trait values are entirely independent across species, meaning there is no phylogenetic correlation or influence of evolutionary history on trait similarity. Under this model, trait values follow a normal distribution with a constant variance, calculated from the trait values, without any shared evolutionary trajectory. The variance-covariance matrix for this model is characterized by non-zero values only on the diagonal, indicating that each species’ trait is unrelated to any others (39).

Another early approach to stepping beyond Brownian motion was suggested by Mark Pagel in 1999 (49). While the originally suggested approach was designed for discrete traits, the author later pointed out that it applies equally well to continuous traits, and has been used extensively ever since. In traditional explicit evolutionary models, such as Brownian motion and its extensions, trait evolution is modeled by explicitly defining and adjusting parameters that represent various evolutionary processes driving trait change. Pagel’s frameworks of tree transformations differ from the traditional explicit evolutionary models discussed above. Instead of directly adjusting the mechanics of the model, Pagel’s tree transformations (λ, δ, and κ) directly adjust the elements of the phylogenetic variance-covariance matrix. This translates into different transformations of branch lengths of the tree.

Pagel’s λ measures phylogenetic signal using the Brownian motion model as a baseline, where values range from 0 (independent trait evolution) to 1 (evolution under Brownian motion), with intermediate values indicating partial phylogenetic influence. The δ transformation adjusts node heights to model changes in evolutionary rates over time, with δ < 1 indicating deceleration and δ > 1 suggesting acceleration toward the present. Finally, the κ transformation alters branch lengths to explore how evolutionary distance affects trait similarity, with κ = 1 retaining the original tree structure and κ = 0 equalizing all branch lengths across the tree, representing a situation where change is proportional to the number of nodes separating species rather than time (37).

### Variations in Evolutionary Mode Across Species

In Table 1, σ^2^ represents one measure of evolutionary rates in terms of the variance in trait evolution. z_0_ denotes the trait value at the root of the phylogenetic tree (ancestral state). t_ij_ refers to the shared evolutionary time between species i and j. α is the strength of stabilizing selection in the Ornstein-Uhlenbeck (OU) model. a is the rate change parameter in the Early Burst (EB) model, where negative values indicate decelerating rates of evolution. λ measures the degree of phylogenetic signal in Pagel’s Lambda model. δ adjusts evolutionary rates as a function of time in Pagel’s Delta model. κ describes evolutionary change following a speciational pattern in Pagel’s Kappa model, scaling the branch lengths. Slope indicates a linear trend in the rate of evolution over time (Rate Trend model), while Drift represents the directional trend in trait evolution over time (Mean Trend model).

**Table 1:**
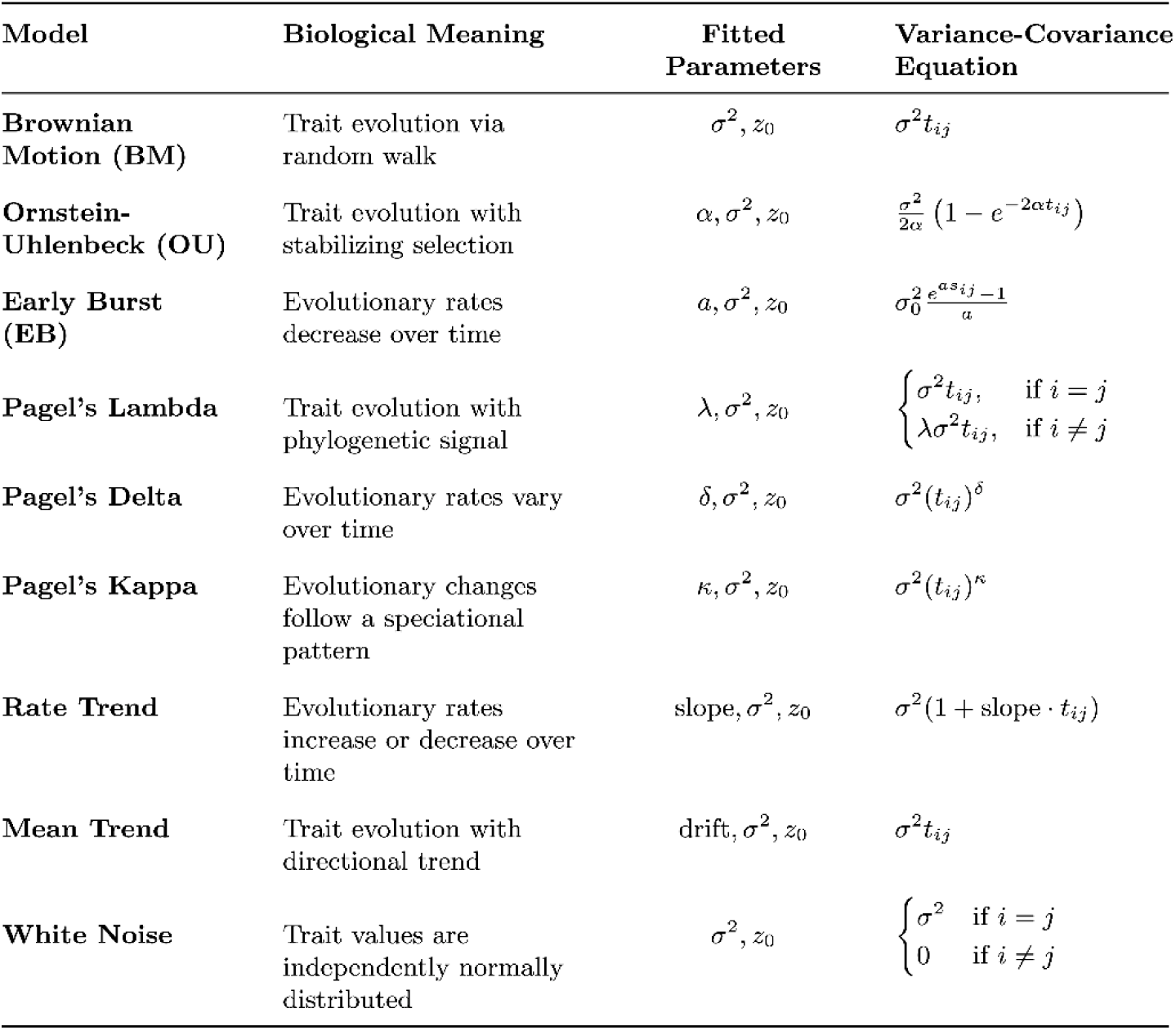
Comparison of models in phylogenetic trait evolution.

### Variations in Best Fit Evolutionary Mode

Across the tree of life, variations in life history strategies, genetics, and environmental pressures have all driven a diverse set of evolutionary solutions for managing disease risk. Instead of assuming a uniform path of evolution, fitting and comparing modes of evolution across groups using *geiger*’s fitContinuous function provides us with a representation of how evolutionary modes can diverge across species. The models are first fit to the overall tree and dataset based on the lowest AIC score, followed by model fitting at the class level. A model with an AIC score at least 10 points lower than the alternatives is considered a significantly better fit. If multiple models fall within 10 AIC points of each other, the evidence for selecting a single best model is weak. In this case the best-fit model from the higher taxonomic classification is designated as the best fit model for the group. This approach maintains consistency in model assumptions, ensuring that for a group to diverge from the evolutionary patterns of its ancestors, it must demonstrate a significant deviation in model fit. See Table 1 for the biological meaning of each evolutionary model.

### Reconstructing Disease Traits

To properly reconstruct the evolutionary rates and disease states for a group of species, the phylogenetic tree and disease data is first fit to various models of evolution to assess the model of best fit. In the case of the mammal data, Pagel’s λ tree transformation is the best fit. The λ value is fit to the phylogeny using the *fitContinuous* function from geiger. This lambda is then used to rescale the phylogeny using the *rescale* function (e.g., λ=0.46), adjusting the branch lengths to reflect the phylogenetic signal in the trait evolution. Ancestral states and confidence intervals under a Brownian motion model for malignancy prevalence were estimated using maximum likelihood, implemented via the anc.ML function from *phytools*. Estimated ancestral states under Pagel’s model are represented by the white line, illustrating the inferred malignancy prevalence at internal nodes of the phylogeny. The surrounding blue haze represents the 95% confidence interval, capturing the range of uncertainty in these reconstructions and highlighting variability in trait evolution across mammalian lineages.

### Adapting Phylogenetic Generalized Least Squares

Through MLE, we found Pagel’s Lambda to be the overall best fit model for the cross-species cancer prevalence dataset. Two other Pagel models; Ornstein-Uhlenbeck and Rate Trend, were the best fit models for smaller clades. The method, *PGLSSeyPagel*, used in Compton et al. to test for statistical relationships between life history and cancer data, assumed Brownian motion(34). With this, we developed a new function, *PGLSSeyOptim*. This method builds upon the *PGLS.SEy*, from phytools (53), and the *PGLSSeyPagel* method created by Diego Mallo. This function adds the ability to assume Brownian Motion, OU, Pagel’s lambda, delta, kappa, white noise, and rate trend. Parameter estimates within the *PGLSSeyOptim* utilize fitContinuous to ensure continuity between the fitting and the trend estimation process.

### Phylogenetic Comparative Methods Currently utilized within Comparative Oncology

Given the plethora of phenotypic traits and the distinct data formats that they assume (e.g., discrete or continuous) there has been an equally diverse set of phylogenetic comparative methods (PCMs) developed to analyze them. Given a specific phenotypic trait and the type of data collected, a mainstay of comparative phylogenetics has been debates on what is the best process method given the context of the data. For any discipline that is eager to integrate the use of PCMs, it can be daunting to face these debates and determine which method is best suited for their specific purpose. Across the field, a standard for testing the relationships between life history and disease traits across species has yet to be set. To understand which method is most appropriate in the context of evolutionary medicine, simulation studies were conducted. In our simulation, we compared two methods used in previous studies alongside our newly created *PGLSSEyOptim* function. The first method, used in Compton et al.’s *Cancer Prevalence Across Vertebrates* is the *PGLSSEyPagel* function(34). The other method is the *Compar.gee* function from *ape.* The Compar.gee function, adapted from the approach described by Paradis and Claude (2002), employs a generalized estimation equation alongside a phylogenetic correlation structure in order to generate log-odd relationships (54). This method does not use a continuous variable, such as species’ level cancer prevalence, but relies upon the occurrences and non-occurrences of the independent variable within each species. This method is used within Bulls et al.’s similar analysis of cancer across tetrapods to perform a binomial regression (33).

### Validating Popular PCMs in Evolutionary Medicine

Within the recent attempts to apply PCMs to cross-species disease prevalence data, there are several statistical methods being utilized for species level trend estimation. To understand which method is best for estimating trends in disease risk across species, we conducted a simulation study. The study used previously collected life history data from PanTHERIA and Amniote along with simulated cancer prevalence data, to compare error rates between multiple phylogenetic statistical methods. (50–52). The simulation has two changing parameters, which are the minimum number of records per species and the degree of the relationship between simulated life history and neoplasia prevalence values (slope). For each of these combinations, with each method, 1000 iterations were completed before moving on to the next combination of parameters. Each iteration of the simulation first randomly samples the needed amount of species from the dataset. From there, the number of records are simulated using the negative binomial distribution. The negative binomial distribution most similarly mirrored the real distribution of records per species. The phylogenetic trees are then cut and standard errors are calculated for the PGLS functions. The relationships are then simulated using the slope-intercept formula, with a small amount of noise to avoid convergence. For simulations where slope is equal to 0, noise is maintained to ensure a non-significant relationship. For compar.gee, Neoplasia Occurrences and Non-Occurrences are generated from the same data used for the PGLS tests.

Within the simulation study, three different models of evolution, OU, BM, and Pagel’s Lambda, were used to simulate trait data with a trend. The trend was randomly chosen for each trial between −2 and 2. For the random noise experiment, both independent and dependent variables were generated randomly. The number of records per species was chosen randomly for each species from the negative binomial distribution with a minimum threshold of 1, 5, 10 or 20 records per species, to mimic different studies in the literature(17,31,32,34). For the experiments with a given trend and model, a false negative error was indicated if the p-value was greater than 0.05. For the null experiment, false positive errors were indicated if the p-value was less than 0.05. Across all parameter values and methods, the average error rates were 15.9% for the compar.gee binomial, 6.9% for PGLSSey, and 5.84% for PGLSSey.Optim. Additional summary statistics are in supplementary table 2.

## Discussion

### Modes of Evolution Between Clades

Figure 1 illustrates how the best-fit evolutionary models can vary across different branches of a phylogeny. While a single model may explain trait evolution over time across a large group of species, closer examination of specific clades reveals shifts in these models. Under the life history framework, natural selection leads species to adopt different trade-offs to optimize survival and reproduction. As species diverge and follow distinct life history strategies, their evolutionary trajectories can vary as a result. These changes in trait evolution often reflect shifts in those evolutionary trajectories, as species adapt to different ecological pressures and life history demands (20,55–57). Applying a single model across hundreds of species may fail to capture the diverse dynamics of trait evolution, especially when these species experience different evolutionary pressures. Utilizing multiple evolutionary models better reflects the varying rates and processes of evolution within clades. This approach provides a more accurate depiction of the complex, non-uniform evolutionary paths that shape adaptations, including those related to disease prevention. As Beaulieu et al. (2012) argue, biological reality is too complex for any single model to fully capture, and the fit of a model depends on both its parameters and the available data (58). Using multiple models helps account for the varied evolutionary pressures across species.

**Figure 1.**
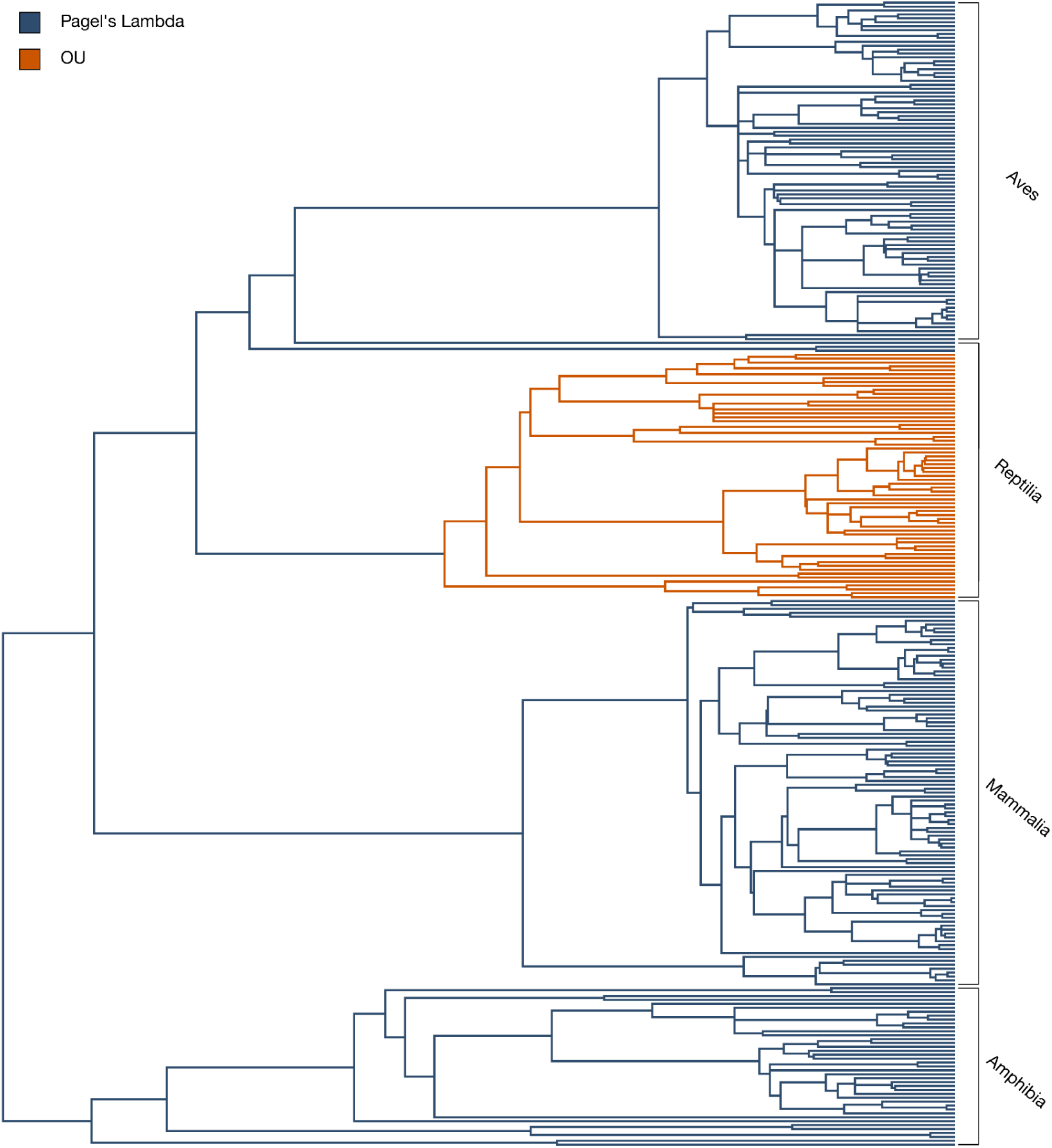
One phylogeny, multiple modes of evolution.

**Figure 2.**
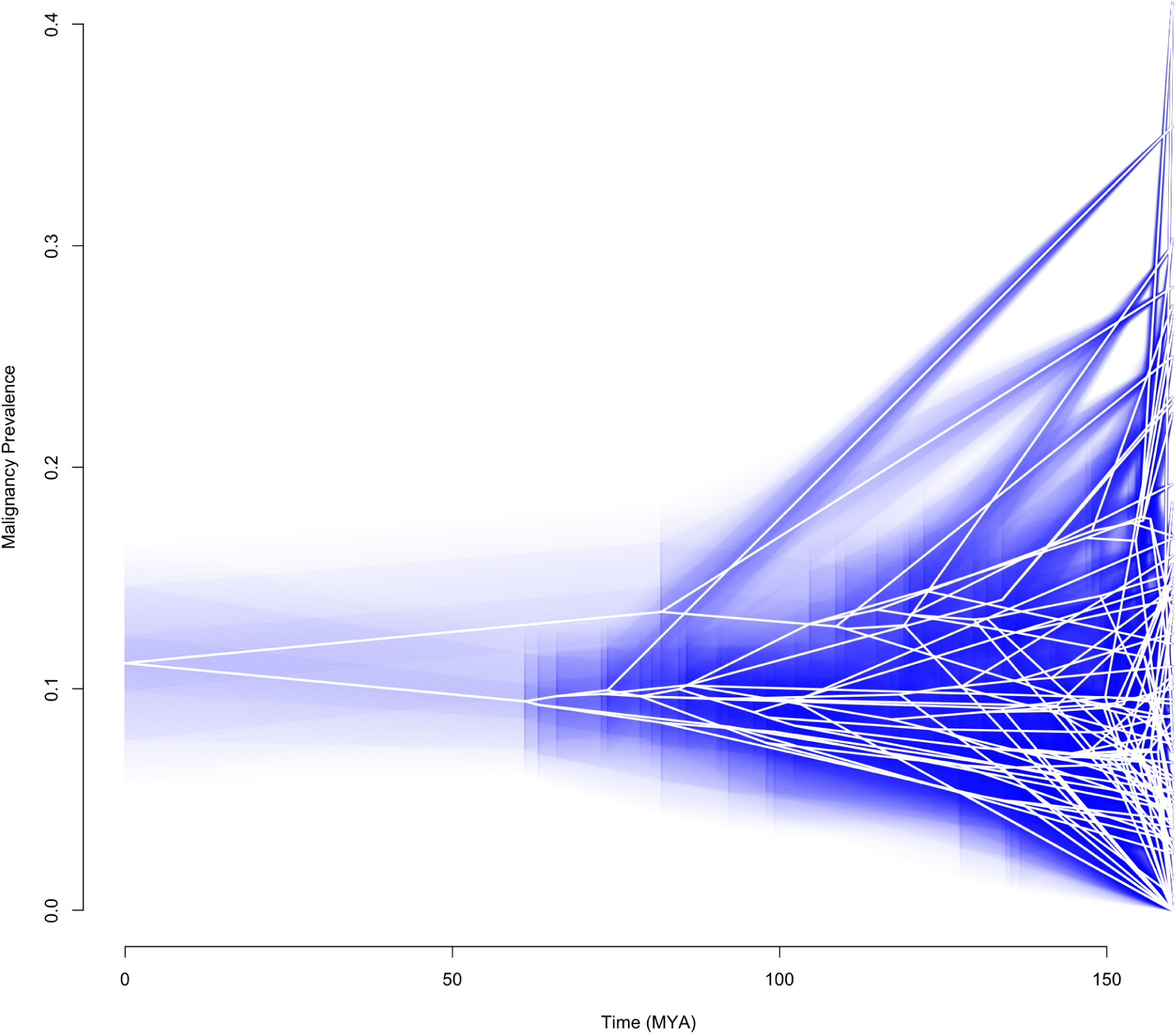
Ancestral State Reconstruction of Malignancy Prevalence under Pagel’s Lambda Model Over Time for 98 Mammalian Species.

### Insights from Ancestral State Reconstruction

Understanding the evolutionary history of cancer prevalence across species has significant implications for human medicine. Evolutionary biologists can utilize ancestral state reconstruction under a best-fit model of evolution to infer how disease traits have changed over time across lineages. If specific genetic variations or evolutionary pressures are associated with higher or lower cancer risk in certain lineages, these findings can guide human cancer research by highlighting conserved or divergent biological pathways involved in tumor suppression, DNA repair, or immune response (59–63). Ancestral state reconstructions can be applied beyond cancer research, providing valuable insights into the evolutionary drivers of various diseases that impact human health.

### Best Practices for Evolutionary Analyses

Our analyses indicate that the PGLS approach provides greater accuracy in detecting trends compared to the binomial regression (Fig 3, ST2). The robustness and additional functionality of PGLSSeyOptim, in conjunction with first fitting for the best evolutionary model for trait data, provides a rigorous framework to ensure that the most appropriate assumptions are made when testing for relationships across a phylogeny. This approach parallels that of Brocklehurst in 2016, who similarly tested for the best fit evolutionary model to analyze evolution rates and variations of body size (64). As phylogenetic comparative methods (PCMs) evolve and become central tools in evolutionary medicine, it is critical to account for the different evolutionary processes that may vary for different traits and trees. Various disease traits may follow distinct evolutionary modes. Failing to account for potential variations in evolutionary models could lead to misinterpretations of phylogenetic relationships and trait correlations. When utilizing tools like PGLSSeyOptim, users should seek the best fit evolutionary model for their data and phylogeny, then utilize the best fit model and its estimated parameters for further phylogenetic analysis.

**Figure 3.**
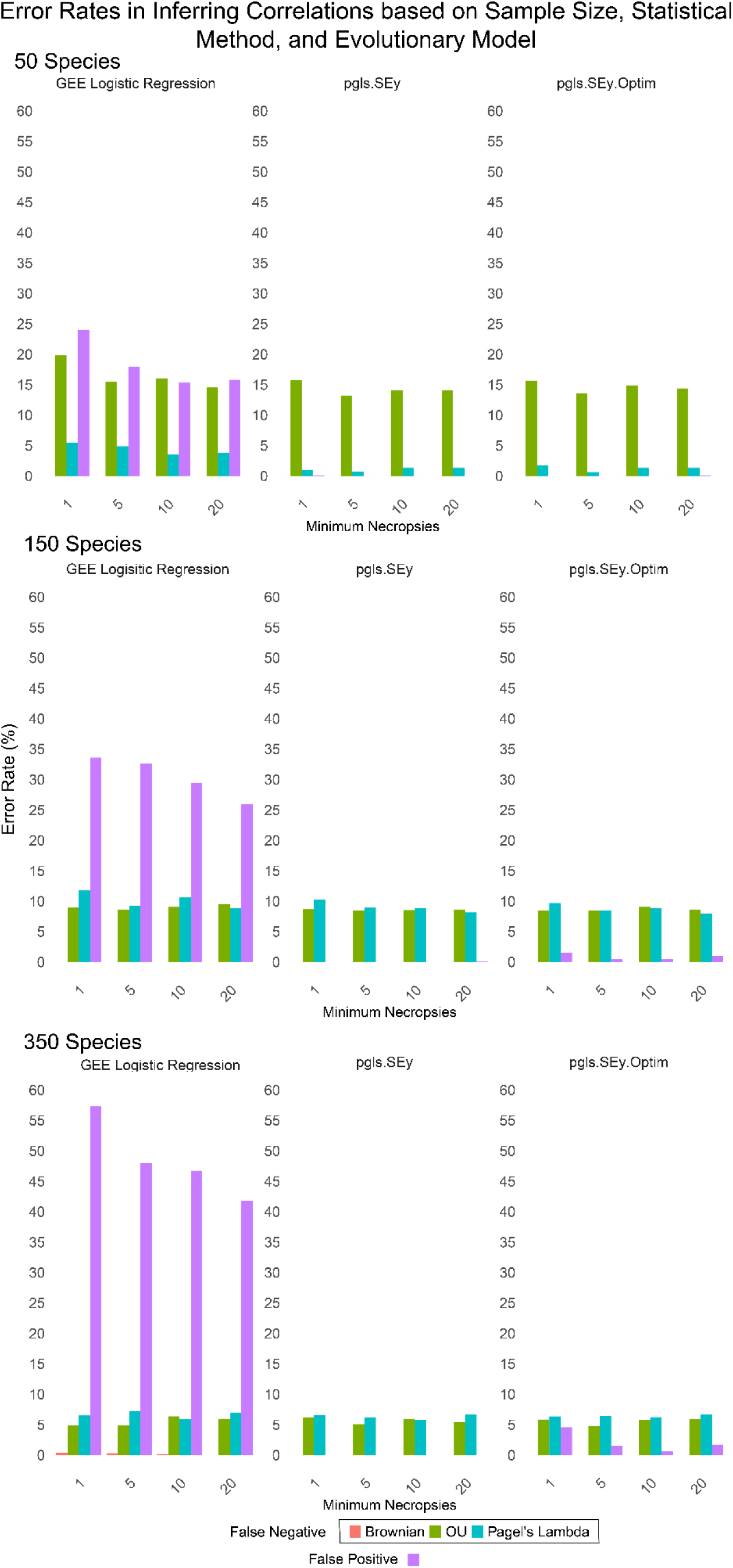
Simulation results for minimums of 1,5,10, and 20 records for 50, 150 or 350 random mammalian species. False positive error rates were based on simulations with no relationship between the independent and dependent variables. False negative error rates were calculated based on simulated data that had a linear relationship between the independent and dependent variables with a slope randomly chosen uniformly between −2 and 2, but that changed along the phylogeny based on one of three models: Brownian motion (Brownian), Ornstein-Ulenbeck (OU) or Pagel’s Lambda.

### Comparison of Phylogenetic Regression Models

All regression methods had similar false negative rates, across modes of evolution, number of species, and number of observations per species (Figure 3). They all appear to be well suited for analyzing disease susceptibility if it evolved randomly over evolutionary time (Brownian motion). That is, there were very few instances of false negative errors when we simulated the trait evolving via Brownian motion, regardless of the type of regression used. However, when disease susceptibility evolved under the OU model, with sudden discrete changes, all regression methods were susceptible to some level of false negatives, which was worse when there were fewer species in the analyses. The standard PGLS method (*pgls.SEy*) performed similarly to our optimized version (*pgls.SEy.Optim*), which estimates parameters within the evolutionary model used to fit the data. With just 50 species in the analysis, both PGLS regressions worked surprisingly well on data generated with the Pagel’s Lambda model, resulting in very few false negative errors. We did find that the binomial regression models are vulnerable to detecting false positive results, and this actually gets worse with more species added to the analyses (Figure 3). This discrepancy likely stems from how the underlying distribution of the necropsy data does not align well with the binomial model’s assumptions, leading to overdispersion. Overdispersion occurs here as the binomial model underestimates the variance inherent in the data, resulting in artificially low standard errors and inflated test statistics. Consequently, binomial regression models appear to fit the data better than they truly do, increasing the risk of false positives. The nature of zoological databases will likely result in a distribution of records that follows a negative binomial distribution. If using count data, researchers should ensure that their data follows the expected distribution of their given statistical method. The MCMCglmm R package allows for count data input following a negative binomial distribution(65). Other groups have employed regressions using phylogenetic independent contrasts instead of phylogenetic VCV matrices to mitigate potential errors in regression estimates caused by phylogenetic correlations. This approach is particularly useful when researchers aim to minimize the influence of phylogenetic signal and evolutionary history in their analyses (66). However, when studying evolving disease traits in a comparative setting, incorporating phylogenetic relationships is crucial for understanding trends within an evolutionary context.

Comparative phylogenetics has a clear, complementary place alongside comparative genomics in explaining the origin, risk, and persistence of human pathologies. As comparative data on diseases in non-human species increases, so shall the need for the robust utilization of comparative phylogenetic methods. Like many other subfields within evolutionary medicine, comparative oncology is a nascent field and we should expect to see an array of methodological approaches to other diseases as well (e.g., studying cardiovascular disease across the tree of life). However, just as genomics has managed to do, there must be agreed upon “best practices” for the analyses of disease prevalence data. While these best practices must be an emergent property of peer-review and discussion amongst the field, we hope that we have at least laid the groundwork for how to evaluate the robustness of these methods within evolutionary medicine.

## Methods

### Aggregating Necropsy Data for Species Level Analysis

We utilized a previously published dataset of cross-species cancer prevalence, containing data on 292 species and 16049 individuals (34). In short, the database is built on the necropsy records of each individual, collected, with permission, from 99 zoological institutions, aquariums, and animal housing facilities. The individual necropsy data was then aggregated, where relevant variables such as benign and malignant tumor incidence were summarized to represent each species. Species level data also includes the number of necropsy records and taxonomic information. The summarized necropsy data was then paired with species-specific adult mass data. Life history trait data was collected from two of the popular life history databases often utilized in comparative oncology studies: PanTHERIA and Amniote (50–52).

### Fitting Popular Models of Evolution to Cross-Species Cancer Prevalence Data

The fitContinuous function from the ***geiger*** package allows for the fitting of phylogenies and disease trait data to various models of evolution(40). The function utilizes Maximum Likelihood Estimation (MLE) to assess the log-likelihood for each model. It iterates through different parameter estimates using the subplex optimization algorithm to maximize the log-likelihood and generate the best parameter estimates. Before fitting, trait prevalence data undergoes an arcsine square root transformation to ensure it follows a normal distribution. The AIC (Akaike Information Criterion) is then utilized to determine the best model, while accounting for overfitting.We applied this framework to fit evolutionary models to cancer prevalence data from 292 species. Based on AIC scores, Pagel’s Lambda tree transformation provided the best fit (Log-Likelihood: 124.72; AIC: −243.45; ST1). The model was fit with an intermediate lambda value of 0.46. This is typically described as an intermediate phylogenetic signal, which indicates that species tend to have similar traits as their close relatives, but not to the extent predicted by Brownian motion.

## Conclusion

Evolutionary medicine has made important insights into the origin and persistence of human disease. As the field has diversified, it has continued to permeate new fields of disease and demonstrated its broad utility to medicine. Despite its successes, much of the progress made in evolutionary medicine has been conceptual, expanding our understanding of human disease within the already established framework of what we know about humans’ evolutionary history. Primary explorations of human-relevant disease are possible through the classical methods employed in evolutionary biology, namely comparative phylogenetics. When we compare human diseases across species the prevalence of that disease is well understood as a continuous species trait. Virtually every human disease that can be viewed through the lens of epidemiology (i.e., a measurement of its prevalence in a population) can be studied in a similar way across species through comparative phylogenetics. As comparative phylogenetics becomes increasingly integrated into disease research, it is essential to establish best practices to ensure accurate and meaningful results that will significantly contribute to our understanding of cancer.

## Supplementary Materials

All code, tables, and figures are available at https://github.com/wjmellon/EvMedPCMS

